## supplementary information for "Transmissibility of clinically relevant atovaquone-resistant *Plasmodium falciparum* by anopheline mosquitoes"

### Supporting Information for
Transmissibility of clinically relevant atovaquone-resistant *Plasmodium falciparum* by anopheline mosquitoes

Victoria A. Balta[a,b], Deborah Stiffler[a,b], Abeer Sayeed[a,b], Abhai K. Tripathi[a,b], Rubayet Elahi[a,b], Godfree Mlambo[a,b], Rahul P. Bakshi[b,c], Amanda G. Dziedzic[a], Anne E. Jedlicka[a], Elizabeth Nenortas[c], Keyla Romero-Rodriguez[c], Matthew A. Canonizado[c], Alexis Mann[a,b], Andrew Owen[d], David J. Sullivan[a,b], Sean T. Prigge[a,b], Photini Sinnis[a,b], Theresa A. Shapiro[a,b,c,1]

Corresponding author: Theresa A. Shapiro

**This PDF file includes:**

Figures S1 to S9
SI Reference

A

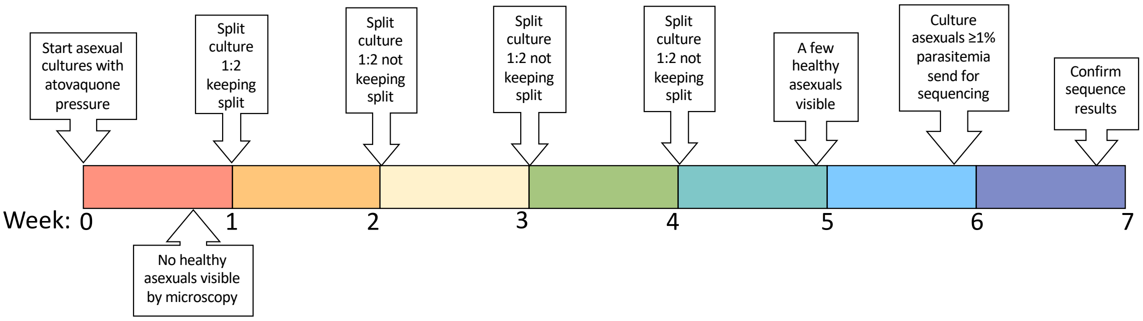

B

| | Parent Strain | Starting Culture Volume (ml) | Hematocrit | Total # of RBCs | Starting Parasitemia | Total # parasites pressured | Atovaquone concentration (nM) | Successful isolates | Mutant cells first evident in culture |
|---|---|---|---|---|---|---|---|---|---|
| **Trial 1** | NF54 | 100 | 4% | $2.20x10^{10}$ | 3.0% | $6.60x10^{8}$ | 70 | Y268S | day 34 |
| **Trial 2** | NF54 | 100 | 4% | $2.20x10^{10}$ | 3.0% | $6.60x10^{8}$ | 35 | L144S | day 29 |
| **Trial 3** | NF54 | 100 | 4% | $2.20x10^{10}$ | 3.0% | $6.60x10^{8}$ | 20 | Y268S | day 25 |
| **Trial 4** | NF54 | 50 | 4% | $1.10x10^{10}$ | 3.0% | $3.30x10^{8}$ | 70 | None | |
| **Trial 5** | NF54 | 100 | 2% | $1.10x10^{10}$ | 4.0% | $4.40x10^{8}$ | 70 | None | |
| **Trial 6** | NF54 | 100 | 2% | $1.10x10^{10}$ | 4.5% | $4.95x10^{8}$ | 70 | None | |
| **Trial 7** | NF54 | 50 | 4% | $1.10x10^{10}$ | 3.0% | $3.30x10^{8}$ | 35 | None | |
| **Trial 8** | NF54 | 100 | 2% | $1.10x10^{10}$ | 3.0% | $3.30x10^{8}$ | 10 | L144S | day 47 |
| **Trial 9** | NF54 | 100 | 2% | $1.10x10^{10}$ | 5.0% | $5.50x10^{8}$ | 50 | None | |
| **Trial 10** | NF54 C21 | 50 | 2% | $5.50x10^{9}$ | 7.4% | $4.07x10^{8}$ | 70 | None | |
| **Trial 11** | NF54 C21 | 50 | 2% | $5.50x10^{9}$ | 6.0% | $3.30x10^{8}$ | 35 | Y268S | day 35 |
| **Trial 12** | NF54 C3 | 100 | 4% | $2.20x10^{9}$ | 6.0% | $1.32x10^{9}$ | 70 | none | |
| **Trial 13** | NF54 C19 | 100 | 4% | $2.20x10^{9}$ | 6.0% | $1.32x10^{9}$ | 70 | none | |

C

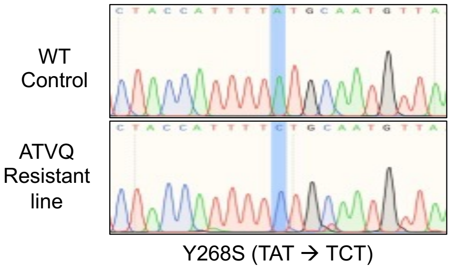

**Figure S1.** In vitro selection of gametocyte-competent, atovaquone-resistant, *P. falciparum*. (A) Representative timeline for selecting atovaquone-resistant parasites from under constant drug pressure. (B) Compilation of thirteen attempted selections. NF54 parent line was known to generate infectious gametocytes; parasites from trials 10-13 were low passage isolates from clinical trial volunteers (1). Total number of parasites pressured was 7.8 x $10^{9}$. (C) Sequence data revealing an A-to-C mutation resulting in Y268S change. No evidence for heteroplasmy was detected in any of the WT or mutant samples.

## A

| Primer Code | Sequence (5' to 3')[a] | Product (bp) | Primer Description |
|---|---|---|---|
| **Full length sequence of *Pfcytb* from cells that survive atovaquone pressure** | | | |
| CytBSeqF | GGTAATGCTGCCATTGATGTAGCATTAC | 1848 | Forward for amplification |
| CytBSeqR | GCATGCAATACCGAACATTTATCG | | Reverse for amplification |
| CytBSeqF1 | CGTTGGTTATGTCTTACCATGGGG | - | For sequencing |
| CytBSeqF2 | GGAATTATACCTTTATCACATCCTGATAATGC | - | For sequencing |
| **Nested PCR of *Pfcytb* to detect parasite DNA in tissues** | | | |
| CytB1 | CTCTATTAATTTAGTTAAAGCACA | 939 | Forward for primary PCR |
| CytB2 | ACAGAATAATCTCTAGCACC | | Reverse for primary PCR |
| CytB1.1 | CGTTGGTTATGTCTTACCATGGGG | 538 | Forward for nested PCR |
| CytB2.2 | AGTTGTTAAACTTCTTTGTTCTGC | | Reverse for nested PCR |
| **RFLP to detect WT and mutant parasites in mixed infections** | | | |
| CytB1 | CTCTATTAATTTAGTTAAAGCACA | 939 | Forward for primary PCR |
| CytB2 | ACAGAATAATCTCTAGCACC | | Reverse for primary PCR |
| CytB5 | GGTTTACTTGGAACAGTTTTTAACAaTG | 250 | Reverse for secondary PCR of WT |
| CytB8 | GTAGCACAAATCCTTTAGGGTATGA | | Forward for both secondary PCRs |
| CytB9 | GGTTTACTTGGAACAGTTTTTAACAcTG | | Reverse for secondary PCR of mutant |

[a]Residues in lower case red encode base changes to generate NsiI (a) or PstI (c) endonuclease sites that distinguish mutant from WT at codon 268.

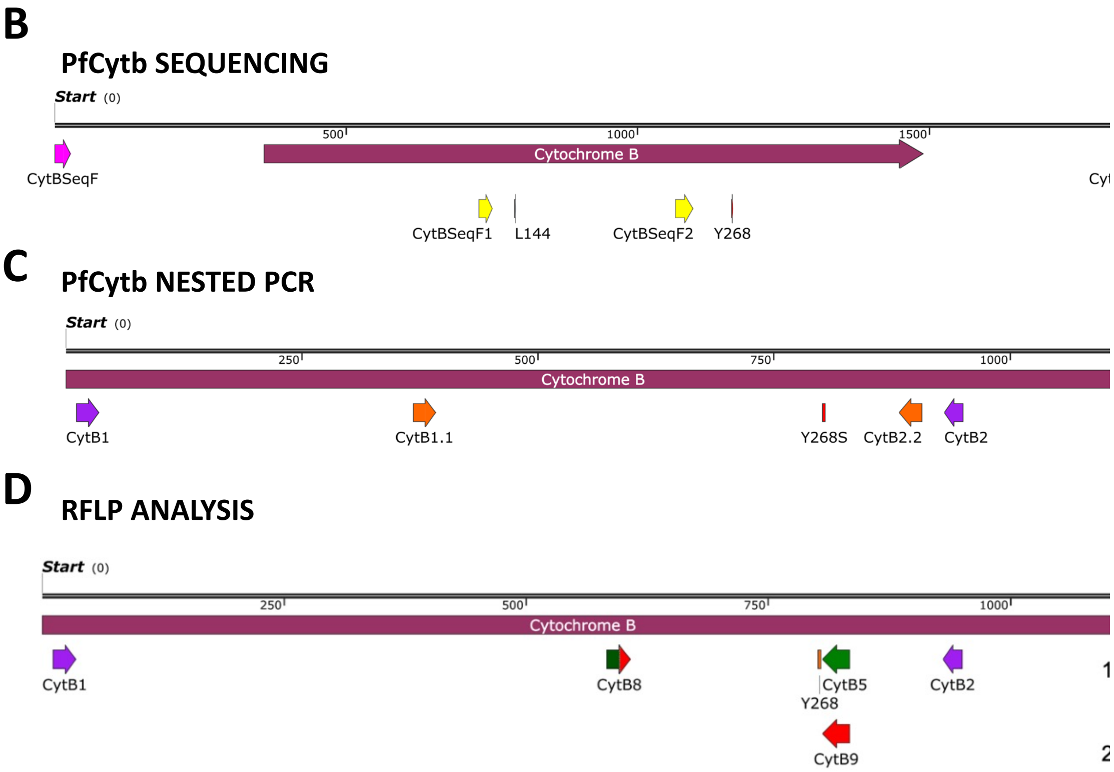

**Figure S2**. PCR primers used in these studies. (A) Primer sequences and descriptions. (B) Graphic of primers for *Pfcytb* sequencing. (C) Graphic of primers for *Pfcytb* nested

PCR. (D) Graphic of primers for RFLP analysis. To identify low levels of Y268S-encoding DNA in mosquito tissues, an RFLP method (2) was modified as follows. DNA was isolated from midguts and salivary glands (Monarch Genomic DNA purification kit, NEB), and about 10 ng DNA was analyzed. In the primary PCR, a 939 bp product containing the mutation site was amplified with Taq DNA polymerase (NEB). Then 250 bp nested products were amplified by Taq DNA polymerase from 1 μL primary reaction product plus primer pairs: either CytB8 and CytB5 for detection of WT Y268, or CtyB8 and CytB9 for detection of mutant Y268S. Reverse primer CytB5, in conjunction with WT sequence TAT in the template DNA, results in an NsiI recognition site (ATGCAT) in the nested PCR product. Reverse primer CytB9, in conjunction with mutant sequence TCT in the template, generates a PstI recognition sequence (CTGCAG). For RFLP analysis, 5 μL of amplified DNA was digested (37 °C, overnight) with 1U NsiI or PstI (NEB). Products were resolved in 3% agarose and visualized by GelStar (Lonza) fluorescence. Successful digestion by either NsiI or PstI yields 224 and 26 bp products. From mixed cultures, an allelic population as low as 2% (NsiI) or 10% (PstI) can be detected.

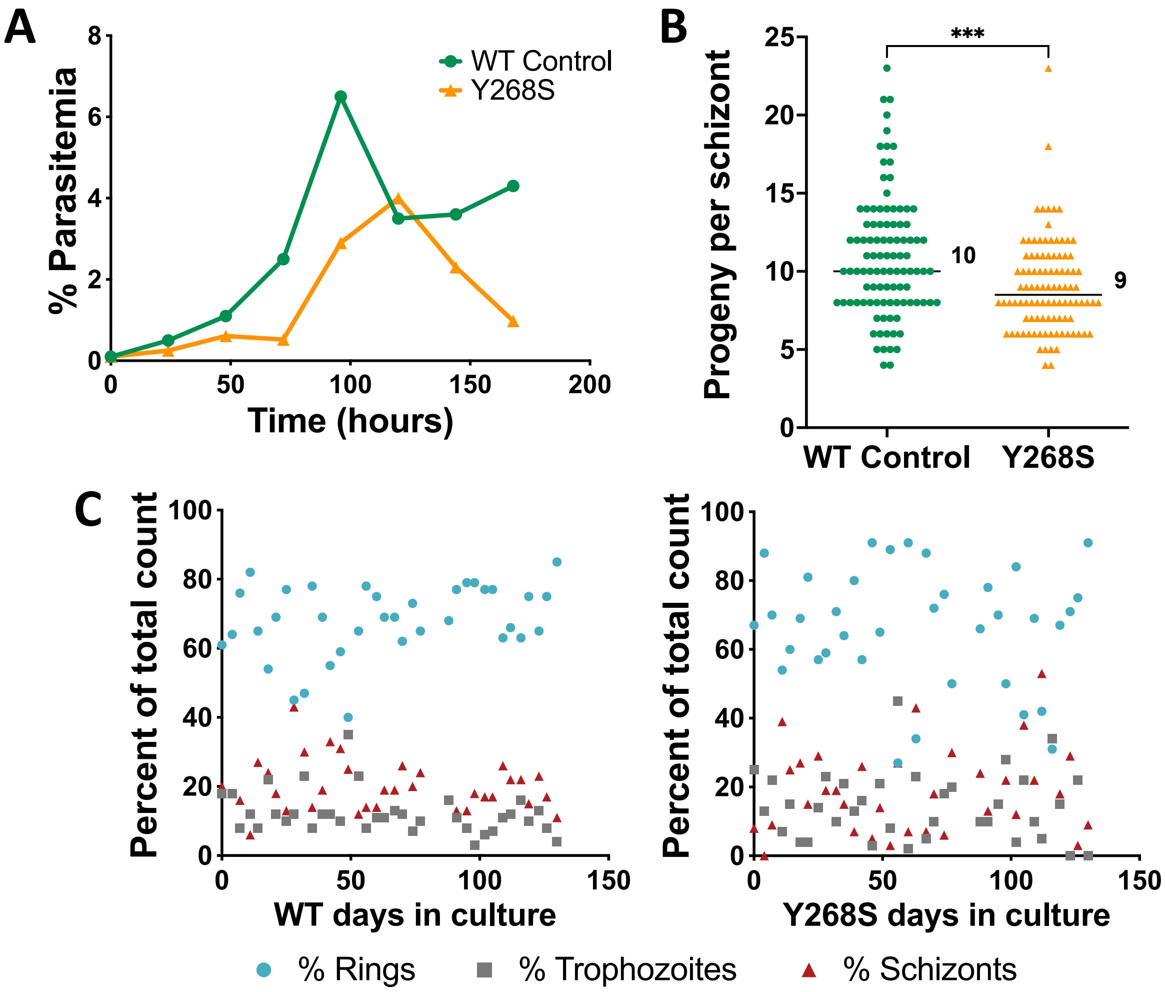

**Fig. S3.** Growth characteristics in vitro of WT and Pfcytb Y268S asexual erythrocytic parasites. (A) Cell count was monitored at indicated intervals for a culture seeded at 0.1% parasitemia and not passaged. Relative to WT, mutant cells had a prolonged lag phase, delayed and lower peak parasitemia, and did not persist in stationary phase. Data from one biological replicate. (B) Progeny within late schizonts were counted in SYBR Gold-stained thin smear by fluorescence microscopy at 1000x magnification. Indicated are median values (ranges 4 to 23 for WT, and 4 to 23 for Y268S); $n$ = 100, ***$P$ = 0.0001. (C) Asynchronous parasites grown continuously over four months and without shaking were sampled for differential count at every 3-4 day passage. Depicted are percent of each stage in WT (*left panel*) and Y268S mutant (*right panel*) cultures. The tendency in WT cultures for trophozoites to be the least abundant form is lost in mutant populations.

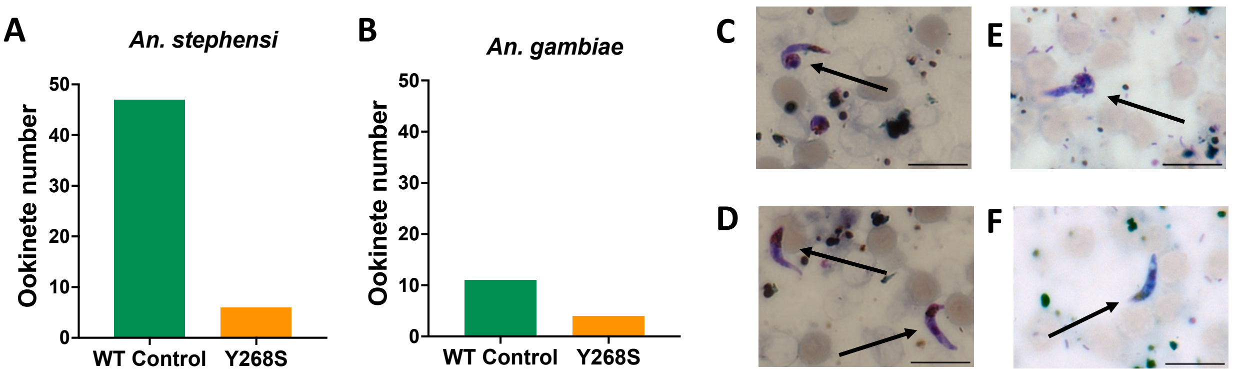

**Fig. S4.** Evaluation of ookinetes in anopheline mosquito midguts at 20-26 h after membrane feed. (A) Sum of mature and immature ookinetes in pooled samples obtained from ten *An. stephensi* midguts, in 60 independent 1000X microscopy fields. Data are from one biological experiment. (B) Sum of mature and immature ookinetes seen in ten *An. gambiae* midguts, in 60 independent 1000X microscopy fields. Data are from one biological experiment. (C) Immature WT control ookinete in *An. stephensi*. (D) Mature WT control ookinetes in *An. stephensi*. (E) Immature WT control ookinete in *An. gambiae*. (F) Mature WT control ookinetes in *An. gambiae*. *Bars*, 10 μm. *Arrows*, representative ookinetes.

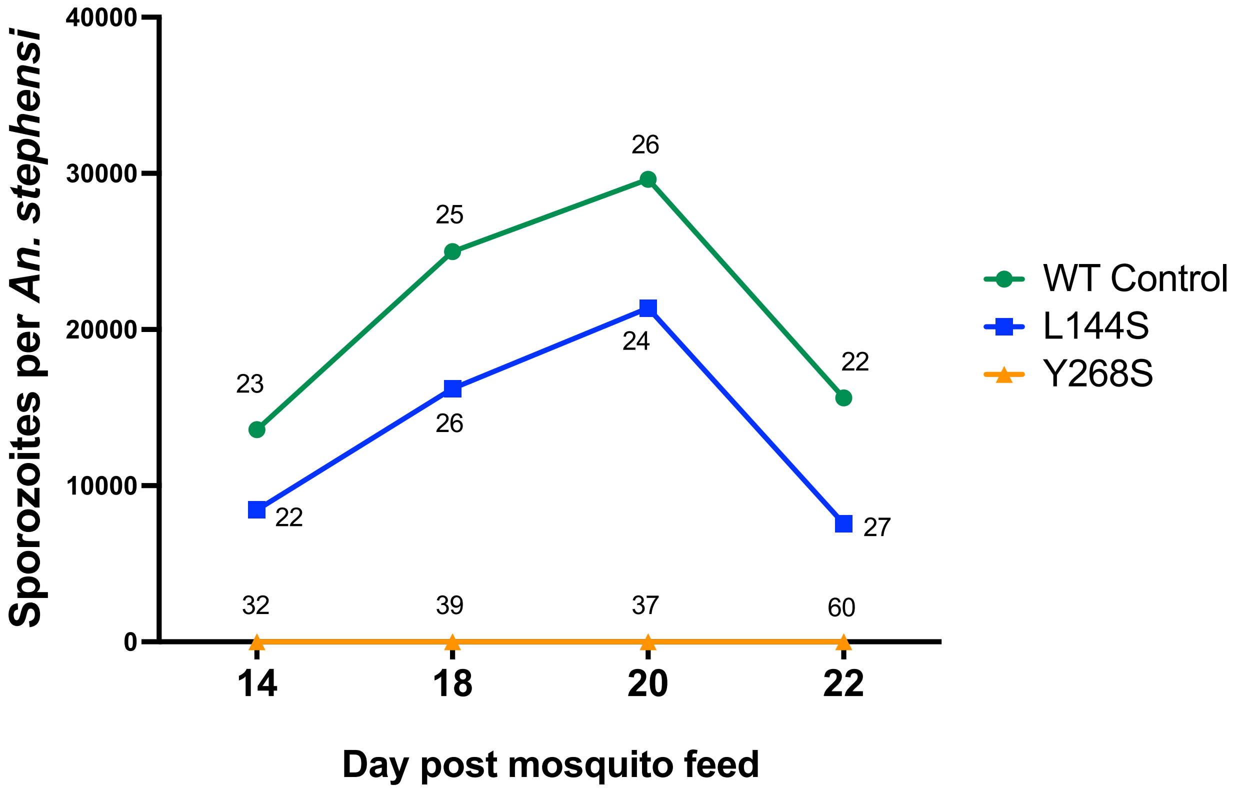

**Fig. S5.** *An. stephensi* salivary gland sporozoite loads over time for wild type or *Pfcytb* mutant *P. falciparum*. At indicated intervals after bloodmeal, salivary glands were harvested and sporozoite load was assessed. Data are from one biological replicate. Number of mosquitoes dissected per timepoint for each experimental group is annotated, totaling 96 for WT, 99 for L144S, and 168 for Y268S.

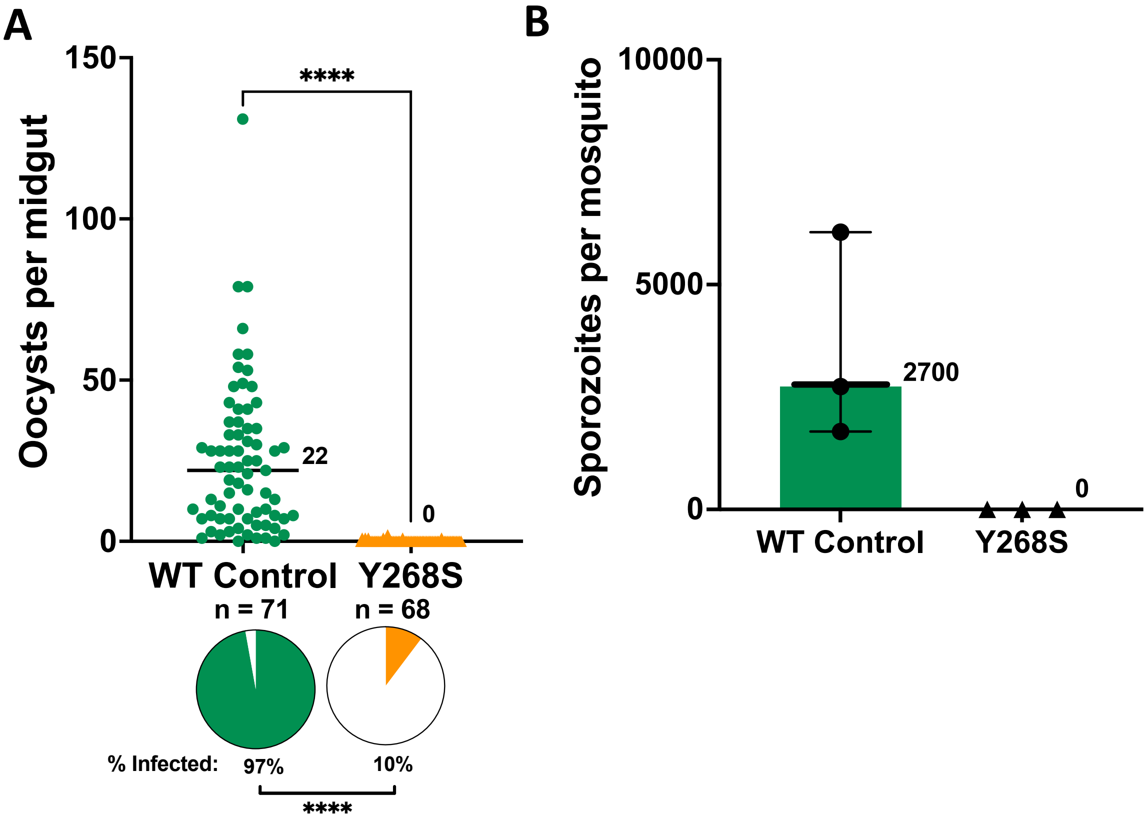

**Fig. S6**. *P. falciparum* counts in tissues of *An. gambiae*, from three biological replicate experiments. (A) Midguts were examined at 7-10 d after membrane feed. Median number of oocysts per midgut is indicated (ranges 0 to 131 for WT, or 0 to 2 for Y268S); *n*, total number of mosquitoes dissected, ****$P$ <0.0001; *pie charts,* percent of mosquitoes infected, ****$P$ <0.0001. (B) Salivary glands were dissected 14-19 d after membrane feed, from a total of 90 mosquitoes infected with WT or 100 infected with Y268S mutant parasites. *Symbols*, for each experiment, average number of sporozoites per mosquito. *Bars*, median values (range 1700 to 6200 for WT, no sporotoites were seen in any Y268S sample); not powered for statistical analysis.

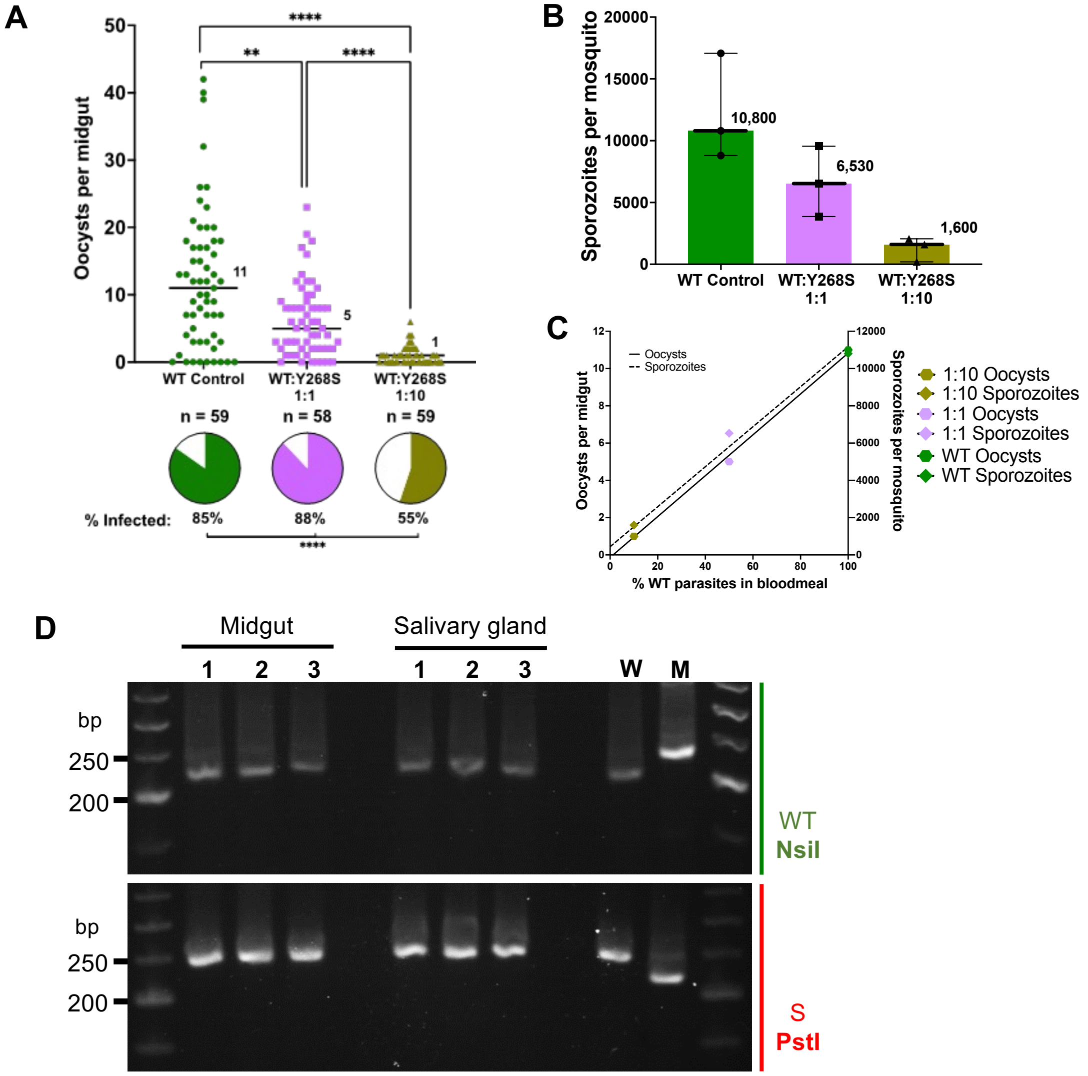

**Fig. S7.** Mixed infection with WT and Y268S mutant *P. falciparum* in *An. stephensi* mosquitoes. Parasites were fed to mosquitoes in three cohorts: WT control (0.5% gametocytemia), a 1:1 mixture of WT:Y268S (0.25% WT:0.25% Y268S gametocytemia), or a 1:10 mixture of WT:Y268S (0.05% WT:0.45% Y268S gametocytemia). Results are from three independent experiments. (A) Oocysts were counted at 7-9 d after membrane feed. *Bars* are median (values indicated); WT range 0 to 42, 1:1 range 0 to 23, 1:10 range 0 to 6; *n*, number of mosquitoes dissected. Overall Chi-square < 0.0001; ***P* < 0.0040, *****P* < 0.0001 by Kruskal-Wallis. *Pie charts* are percent of mosquitoes infected; *****P* <0.0001 by Fisher's exact. (B) Salivary glands were removed for sporozoite counts 14-16 d after infection. Total number of mosquitoes were 90 for WT, 91 for 1:1, and 90 for 1:10. *Symbols*, average sporozoites/mosquito for each experiment; *bars*, median (values indicated; range 8,800 to 17,000 for WT; 3,900 to 9,500 for 1:1; 200 to 2,100 for 1:10). Overall Chi-square 0.039; not powered for further analysis. (C) Parasite numbers as a function of percent WT gametocytes in the bloodmeal. Oocysts per midgut: y = 0.11x – 0.14, $R^2$ 0.998; sporozoites: y = 107x + 440, $R^2$ 0.988. (D) RFLP analysis for Y268S mutant DNA in mosquito tissues. NsiI cleaves only the WT *Pfcytb* sequence (*upper panel*), whereas PstI cleaves only the sequence encoding Y268S (*lower panel*); in both cases a 224 bp product is released (SI Appendix, Fig. S2*A,D*). *1*, WT control; *2*, WT:Y268S at 1:1; *3*, WT:Y268S at 1:10; *W*, WT DNA as assay control; *M*, Y268S DNA as assay control.

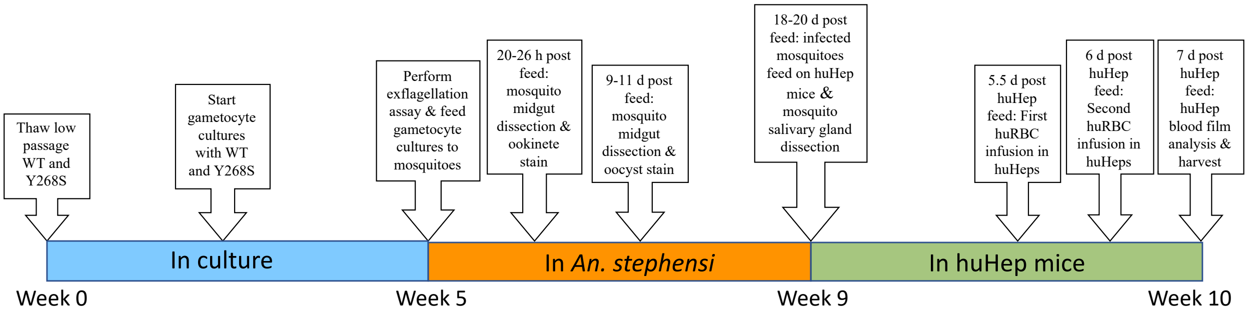

**Fig. S8.** Representative experimental timeline of *P. falciparum* transmission from *An. stephensi* to huHep mice.

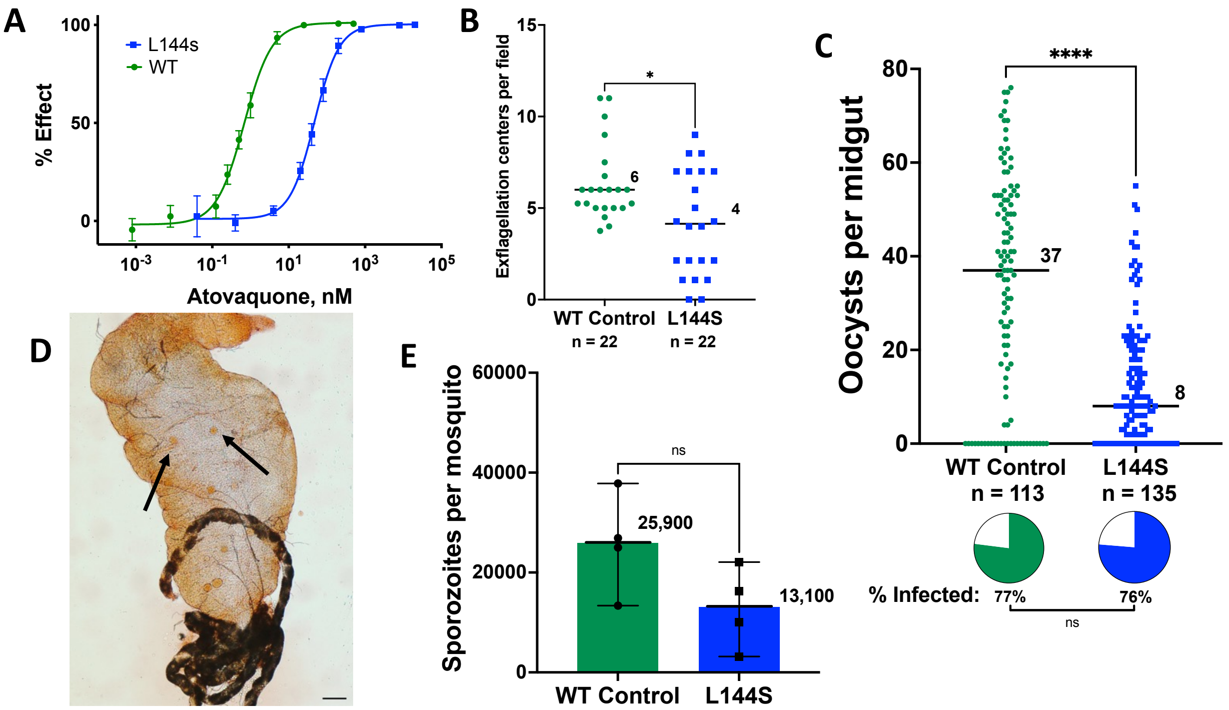

**Fig. S9.** Evaluation of paired wild type and L144S *P. falciparum* in vitro and in *An. stephensi* mosquitoes. (A) Atovaquone cytotoxicity against WT or L144S asexual erythrocytic parasites ($EC_{50}$ 0.68 or 48 nM, respectively). Depicted are mean ± SD of quadruplicate determinations in at least two independent experiments (some SD are too small to extend outside the symbols); $R^2 \geq 0.997$. Susceptibilities of WT and L144S mutant cells to artemisinin (9.8 versus 8.5 nM) or chloroquine (8.2 versus 8.8 nM) were not significantly different. (B) Male gametocyte exflagellation adjusted to 1.5% gametocytemia. Median numbers in two independent biological replicates are indicated (ranges 4 to 11 for WT, and 0 to 9 for mutant); *n*, total number of fields examined; $^*P$ = 0.0253. (C) Oocysts in four independent biological replicates at 9-10 d after membrane feed. Median number of oocysts per midgut is indicated (ranges 0 to 76 for WT, and 0 to 55 for L144S); *n*, total number of mosquitoes dissected; $^{****}P < 0.0001$; *pie charts* percent of midguts infected. (D) Photomicrograph of representative midgut sample from *An. stephensi* infected with L144S *P. falciparum* gametocytes. *Bar*, 100 µm; a*rrow*, L144S oocyst. (E) *P. falciparum* sporozoite counts in four independent biological replicates at 17-20 d after membrane feed, entailing 109 WT-fed and 118 mutant-fed mosquitoes. *Symbols*, for each experiment, average number of sporozoites per mosquito; *bars*, median number of sporozoites (range 13,000 to 38,000 for WT, 3,100 to 22,000 for mutant); *P* = 0.1143.
